# Higher vascularity at infiltrated peripheral edema differentiates proneural glioblastoma subtype

**DOI:** 10.1101/2020.04.17.046466

**Authors:** Eduard Chelebian, Elies Fuster-Garcia, María del Mar Álvarez-Torres, Javier Juan-Albarracín, Juan M. García-Gómez

**Author notes:** Corresponding author: Name: Eduard Chelebian, Full name: Eduard Artur Chelebian Kocharyan.

## Abstract

**BACKGROUND AND PURPOSE:** Genetic classifications are crucial for understanding the heterogeneity of glioblastoma. Recently, MR perfusion imaging techniques have demonstrated their ability to determine molecular alterations. In this work, we investigated whether perfusion markers within infiltrated peripheral edema were associated with proneural, mesenchymal, classical and neural subtypes.

**MATERIALS AND METHODS:** ONCOhabitats open web service was used to obtain the cerebral blood volume at the infiltrated peripheral edema for MRI studies of 50 glioblastoma patients from The Cancer Imaging Archive: TCGA-GBM. ANOVA and Kruskal-Wallis tests were carried out in order to assess the association between vascular features and the subtypes. For assessing specific differences, Mann-Whitney U-test was conducted. Finally, the association of overall survival with molecular and vascular features was assessed using univariate and multivariate Cox models.

**RESULTS:** ANOVA and Kruskal-Wallis tests for the maximum cerebral blood volume at the infiltrated peripheral edema between the four subclasses yielded false discovery rate corrected p-values of <0.001 and 0.02, respectively. This vascular feature was significantly higher (p=0.0043) in proneural patients compared to the rest of the subtypes while conducting Mann-Whitney U-test. The multivariate Cox model pointed to redundant information provided by vascular features at the peripheral edema and proneural subtype when analyzing overall survival.

**CONCLUSIONS:** Higher relative cerebral blood volume at infiltrated peripheral edema is associated with proneural glioblastoma subtype suggesting underlying vascular behavior related to molecular composition in that area.

## 1. INTRODUCTION

In the late years, Central Nervous System tumor classification has shifted from being based on microscopic similarities between cells and their levels of differentiation^1^ to additionally include genetic-based features^2^. This is particularly the case for glioblastoma, where several classifications have been defined: on the one hand, the World Health Organization (WHO) classification which distinguishes between *IDH*-wildtype and *IDH*-mutant glioblastomas^2-4^ and, on the other, the Verhaak classification^5^, consisting of 4 subtypes depending on mutations and molecular profile of various cancer-related genes. These subtypes are the mesenchymal, classical, neural and proneural, the latter being related to *IDH* mutations^5,6^. These new classification paradigms have improved the estimation of prognosis^7,8^ and proposed specific therapeutic targets^9-12^, especially for patients with proneural and mesenchymal type glioblastoma.

Considering that Magnetic Resonance Imaging (MRI) perfusion biomarkers have been associated with patients’ overall survival^13-15^ and cellular features^16,17^, several studies were performed to analyze if there was a relationship between vascular biomarkers and the genomic subtypes classifications. Barajas *et al.* studied the influence of glioblastoma genetic and cellular features over MRI, concluding that they could spot the most malignant regions within the tumor^18^. Jain *et al.* demonstrated that combining Verhaak subtypes with vascularity markers at the enhancing tumor provides additional information as a survival predictor^19^. However, they found that the enhancing and non-enhancing regions of the tumor did not present any significant correlations with the genomic subclassification. Another study proposed that tumor blood volume determined by dynamic susceptibility contrast MR perfusion imaging was related to *EFGR* and to *PTEN* expression in some patients^20^.

Most of these studies focus on the vascularity of the enhancing tumor region and only a few remarked the influence of the non-enhancing part of the tumor including the edema region^18,20^. Gill *et al.*^21^ found molecular differences between Verhaak subtypes performing MRI-localized biopsies in the peripheral edema region. Similarly, Price *et al.*^22^ discovered that metabolic and perfusion changes in this region could be found using multimodal MR images. In this sense, we hypothesize that the vascular parameters in the invasive margins of glioblastoma could be related to characteristic combinations of mutations.

The purpose of this article is to assess the correlation between the vascularity present at the infiltrated peripheral edema habitat at preoperative stage and Verhaak molecular classification. To do so, we propose the use of a multicentrically validated^23^ automatic open service named ONCOhabitats (https://www.oncohabitats.upv.es) proposed by Juan-Albarracín *et al.*^15,24,25^. To ensure the comparability of our study, the analysis was performed on the TCGA-GBM open database^26^, which contains MR images and molecular information. In the end, we found correlation between peripheral edema vascularity and specially the proneural glioblastoma subtype.

## 2. METHODS

### 2.1. Patient Selection

Our study included retrospective patients with glioblastoma from The Cancer Imaging Archive - TCGA-GBM^26^. The database consists of 262 histopathological validated glioblastoma patients, 66 of which had preoperative dynamic susceptibility contrast enhanced T2*-weighted perfusion (DSC) imaging information. Three of them were excluded because they did not have genomic information available.

The remaining 63 belong to two different institutions with the following distribution: 48 in the first and the rest in the second. From the first institution, 6 were excluded because of poor perfusion acquisition, mainly due to having an incomplete field of view in the DSC images. Additionally, 5 were excluded due to post-processing errors when performing DSC quantification. From the second one, only 2 were not considered for having an incomplete FOV.

The final cohort was made up of 50 primary glioblastoma patients who had all underwent tumor resection. Age distribution (mean years [minimum, maximum]) was: 13 females (55.2 years [17, 74]) and 37 males (59.5 years, [17, 81]); overall (58.4, [17, 81]). According to the Verhaak molecular classification^27,28^, the group of patients would be divided into 10 classical, 17 mesenchymal, 11 neural and 12 proneural subtype glioblastomas, attending to the mutations and markers they presented. The cohort clinical data along with the subtype of each subject can be found in the S1 Table, whose complete information is retrieved from the original at the TCGA-GBM website^26^.

### 2.2. DSC Imaging Acquisition

From both institutions only 11 studies were obtained using 3T magnetic resonance imaging machines, all belonging to the first institution. For the rest of them, 1.5T imagers were used.

DSC perfusion MRI was performed during the injection of the gadolinium-based contrast (0.1 mmol/kg) using 95 dynamics for the first institution and 60 dynamics for the second institution of T2*-weighted gradient echo echoplanar images. The repetition time (ms)/echo time (ms)/flip angle (°) for each institution were 1900/40/90 and 2000/54/30 respectively.

### 2.3. Computing vascular habitats

All the cases were processed using the Hemodynamic Tissue Signature service found in the ONCOhabitats platform^24^. It provides a reproducible^23^ and automated methodology to define ROI, based on the vascular properties of the lesion, which enables for a more accurate study of peripheral regions. After preprocessing, the ONCOhabitats service delineates for habitats within the lesion based on unsupervised analysis. Fig 1 depicts the two basic steps for obtaining the vascular habitats. First, segmentation into enhancing tumor (ET) and non-enhancing edema is carried out using morphological MRI, that is, contrast-enhanced T1 (T1-Gd), T2 and Fluid-attenuated inversion recovery (FLAIR). Then, after DSC perfusion quantification into relative cerebral blood volume (rCBV) and flow (rCBF), these vascular maps are used to perform the segmentation into the high angiogenic tumor (HAT), the low angiogenic tumor (LAT), the potentially infiltrated peripheral edema (IPE) and vasogenic peripheral edema (VPE). The first two mainly inside the enhancing region of the tumor whereas the second two mainly in the edema. Detailed explanations of the platform functioning can be found in the original articles^15,24,25^.

**Fig 1.**
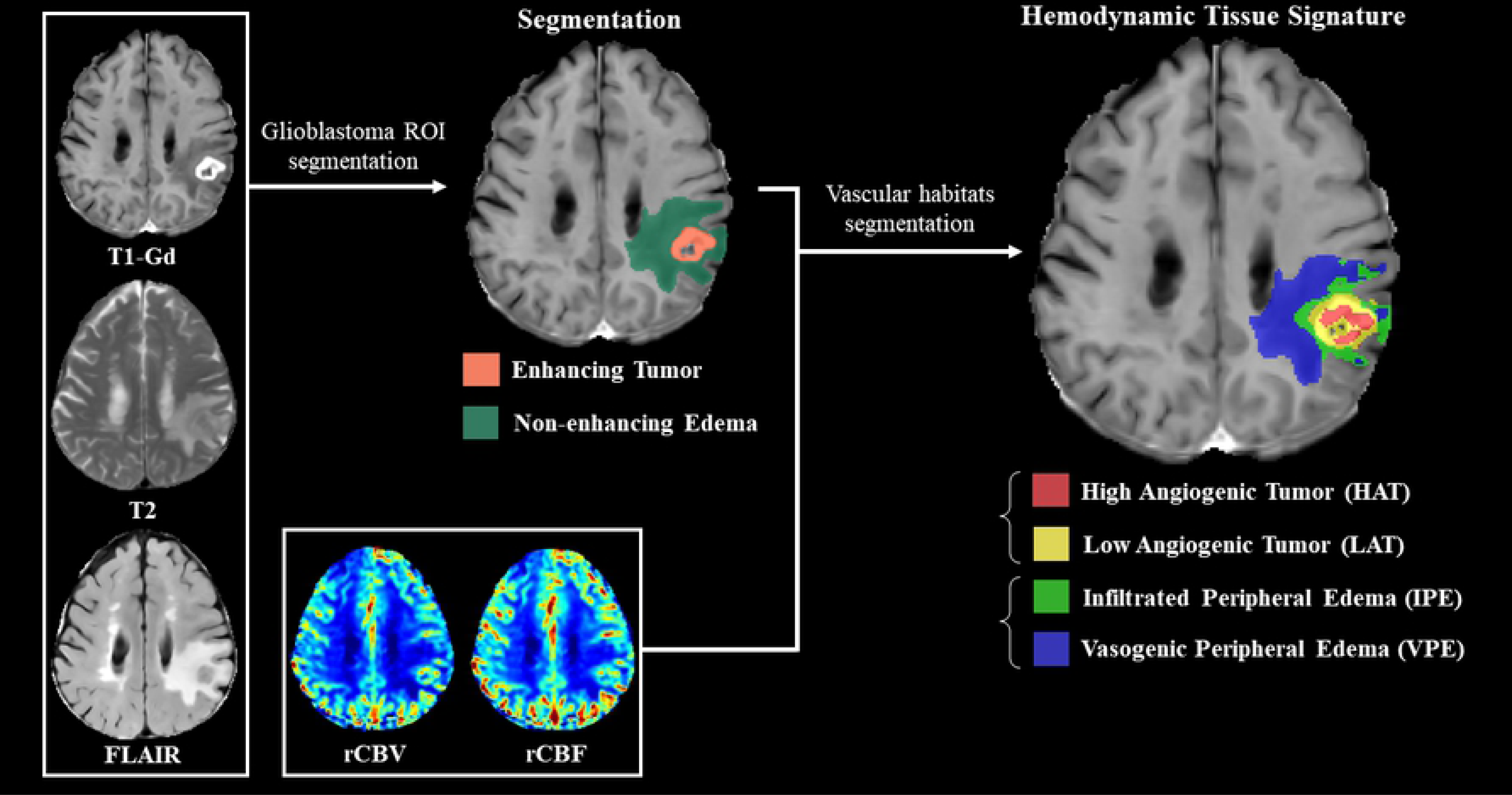
Hemodynamic Tissue Signature pipeline in ONCOhabitats. Morphological sequences are used for segmentation of enhancing tumor and edema. The resulting segmentation together with DSC perfusion maps are used to obtain the vascular habitats. rCBV_max_, rCBV_mean_ and rCBV_median_ were calculated for each vascular habitat. rCBV_max_ was defined as the 95^th^ percentile of the distribution of rCBV values within the ROI in order to increase robustness. Values of rCBV_max_, rCBV_mean_ and rCBV_median_ at each habitat for each subject are presented in the S2 Table.

### 2.4. Statistical Analysis

Firstly, ANOVA and Kruskal-Wallis H test were performed at the ET -in order to have comparable results with the current literature-, consisting of HAT and LAT regions, and at the IPE using all perfusion parameters (rCBV_max_, rCBV_mean_ and rCBV_median_) across Verhaak subclasses. This will serve as a first approximation for establishing any significant divergences in vascularity values regarding the Verhaak subtypes.

For deepening the analysis on the specific differences between the four subclasses, Wilcoxon–Mann–Whitney tests were executed in each habitat. The comparison was made between classical, mesenchymal, neural and proneural classes individually one against the other, and comparing each one of them against the three remaining. For significant experiments, ROC curves were drawn for threshold optimization.

These statistical tests were considered significant when p-values were under 0.05. To correct for multiple testing, Benjamini-Hochberg false discovery rate correction was carried out for every study. Analyses were carried out on a personal computer with MATLAB R2018a (Natick, Massachusetts, USA).

Finally, survival Cox proportional hazards analysis were carried out in order to assess the effect of Verhaak subtypes on Cox overall survival models based only on perfusion parameters. To this end, univariate survival models were fitted using only rCBV_max_ at IPE and at ET. Then, univariate models using only each subtype were fitted. Finally, multivariate models with both rCBV_max_ and each subtype as cofactor were studied. The consequences of subtype addition can be either worsening or improving the fitting, indicating that the subtype provides redundant information or not, that is, there is an association between subtype and vascularity at IPE or not.

For each Cox model, Hazard Ratio (HR) with 95% confidence intervals (CI95), area under ROC curve (AUC) and p-values are reported. Significance will be considered when p-values are under 0.05. Analyses were carried out on a personal computer using R statistical analysis software^29^.

## 3. RESULTS

### 3.1. Verhaak subtypes and rCBV in vascular habitats

In Fig 2 the Box-Whiskers representation of rCBV_max_ values at the vascular habitats for each Verhaak subtype is represented. Firstly, as expected, a decrease in vascularity can be observed as we move further from the central necrotic area (i.e. from HAT to VPE). The proneural subtype shows higher vascularity in every habitat. However, there is important overlap at the enhancing areas, whereas at IPE and, to a lesser degree, at VPE the difference with the rest of the subtypes is bigger. This may point to vascularity differences in the peripheral region.

**Fig 2.**
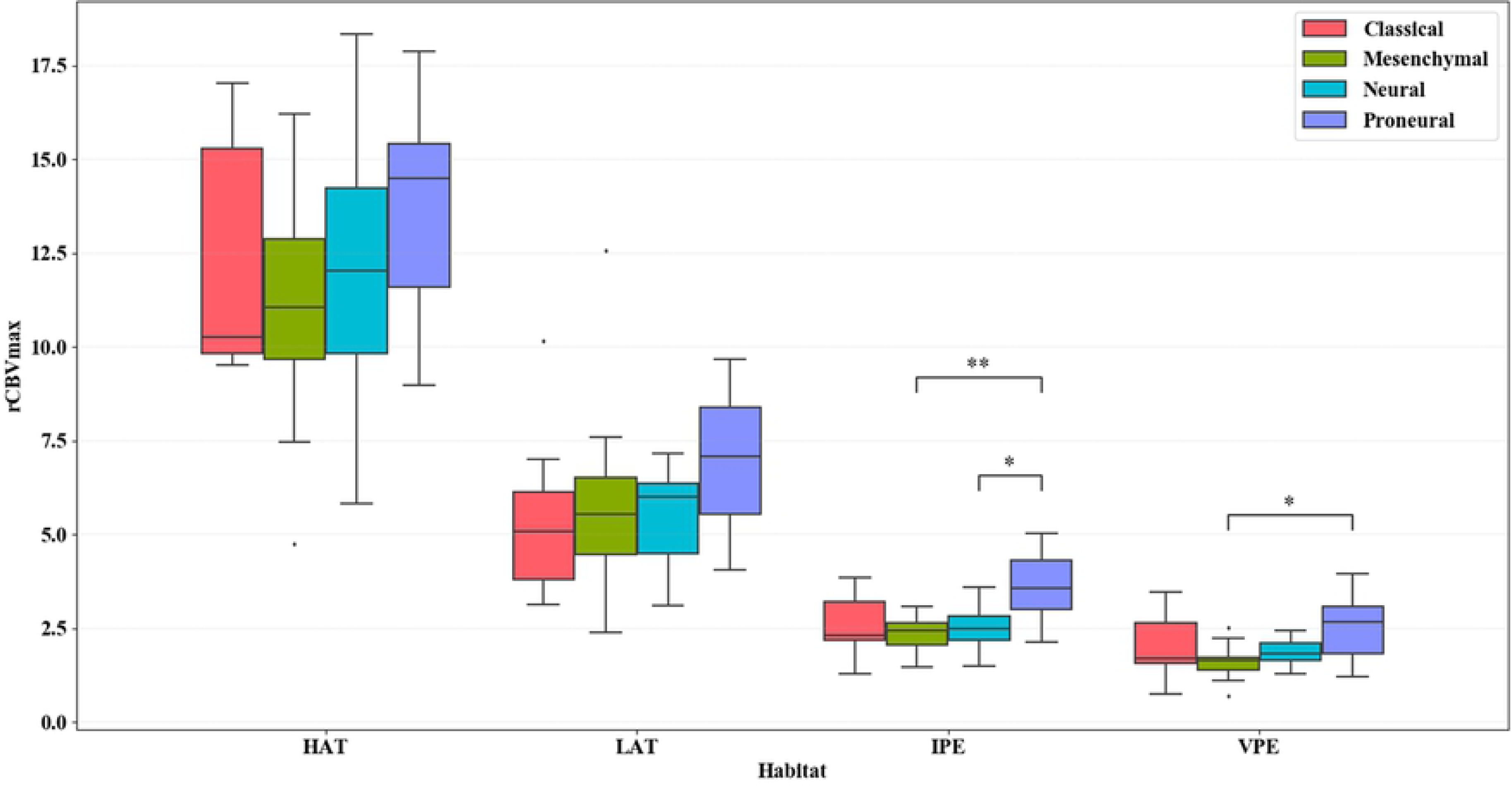
Box-Whiskers representation of rCBV_max_ at each vascular habitat for every Verhaak subtype. Horizontal lines show the significant results of Mann-Whitney tests. * for statistical significance with p<0.05; ** for statistical significance with p<0.01; All p-values are multiple test corrected.

These results are consistent with the ones presented in Table 1. rCBV values at the ET were not significantly different among Verhaak subclasses neither performing an ANOVA nor a Kruskal-Wallis test. However, performing the same analyses on the IPE, significance is found for every rCBV metric (i.e. Max, Mean, Median).

### 3.2. Proneural subtype differences in rCBV at IPE region

Fig 3 shows the mean rCBV at IPE distribution density for each subtype in solid line and dotted lines represent each patient’s rCBV distribution densities at IPE. Subtypes for both mean and individual distributions are represented by different colors. An important difference can be seen in the proneural subtype vascularity distribution, explaining the global differences found in the previous section.

**Table 1.** Mean and standard deviation for rCBV_mean_, rCBV_median_ and rCBV_max_ at ET and at IPE habitat, and p-values from ANOVA and Kruskal-Wallis (K-W) tests in every subtype: classical (Cla), mesenchymal (Mes), neural (Neu) and proneural (Pro).

| rCBV | Region | Cla<br>(n=10) | Mes<br>(n=17) | Neu<br>(n=11) | Pro<br>(n=12) | p<br>ANOVA | p K-W |
| --- | --- | --- | --- | --- | --- | --- | --- |
| **Max** | **ET** | 8.42 ±<br>3.10 | 9.17 ±<br>4.24 | 8.85 ±<br>2.64 | 10.84 ±<br>2.43 | 0.41 | 0.13 |
|  | **IPE** | 2.58 ±<br>0.81 | 2.35 ±<br>0.43 | 2.51 ±<br>0.57 | 3.60 ±<br>0.97 | <0.001* | 0.02* |
| **Mean** | **ET** | 4.01 ±<br>1.74 | 4.14 ±<br>1.42 | 3.98 ±<br>0.92 | 5.27 ±<br>1.40 | 0.11 | 0.11 |
|  | **IPE** | 1.78 ±<br>0.59 | 1.57 ±<br>0.31 | 1.76 ±<br>0.38 | 2.55 ±<br>0.81 | <0.001* | 0.03* |
| **Median** | **ET** | 4.50 ±<br>1.80 | 4.67 ±<br>1.73 | 4.53 ±<br>1.07 | 5.79 ±<br>1.40 | 0.17 | 0.16 |
|  | **IPE** | 1.83 ±<br>0.60 | 1.62 ±<br>0.31 | 1.78 ±<br>0.39 | 2.58 ±<br>0.78 | <0.001* | 0.02* |

All p-values are multiple test corrected; \* for statistical significance.

**Fig 3.**
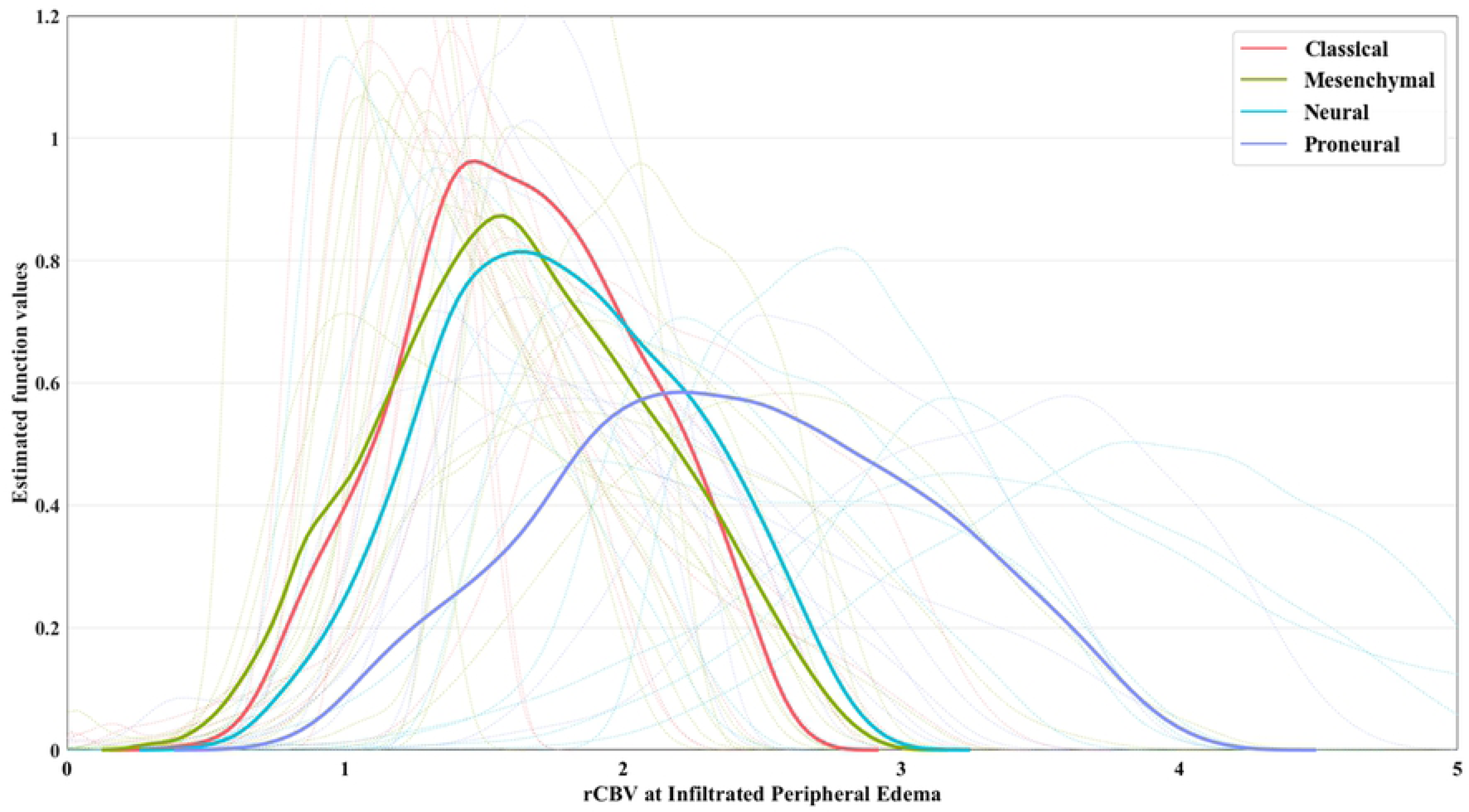
Kernel smoothed density distribution for rCBV values at IPE for each patient and molecular signature. Dotted lines represent density distribution for each patient. Solid lines represent the mean density distribution of patients grouped by molecular subtype.

Results of subtype-specific tests at IPE are presented in Table 2. Comparing rCBV_max_ values at the IPE habitat between the Verhaak subtypes, we obtained that the proneural tumor subtype has a significantly differentiated peritumoral vascularity. Note that the most significant difference was found between proneural and mesenchymal subtype. Mesenchymal subtype vascularity was significantly different from the other three subtypes only before multiple-test correction. Classical and neural subtypes had indistinguishable vascularity at the IPE according to this test, both one from the other and the two from the other subtypes. The same tests were performed for the rest of habitats in the S3 Appendix, yielding only significant results for proneural against mesenchymal rCBV_max_ at VPE.

**Table 2.** Mann Whitney U-test p-values comparing rCBV values at IPE habitat in each subtype against the others individually and rCBV values of each subtype against the rest.

| $rCBV_{\max}$ at IPE | Classical | Mesenchymal | Neural | Proneural |
| --- | --- | --- | --- | --- |
| **Classical** | 1.0000 | - | - | - |
| **Mesenchymal** | 0.9047 | 1.0000 | - | - |
| **Neural** | 1.0000 | 0.7939 | 1.0000 | - |
| **Proneural** | 0.1100<sup>†</sup> | 0.0043<sup>*</sup> | 0.0386<sup>*</sup> | 1.0000 |
| **Rest** | 0.9047 | 0.1023<sup>†</sup> | 0.9047 | 0.0043<sup>*</sup> |

All p-values are multiple test corrected; \* for statistical significance after multiple test correction; † for statistical significance before multiple test correction.

When comparing proneural rCBV_mean_ and rCBV_median_ at IPE against the rest of the subtypes, significant differences were also found (corrected p-values of 0.0428 and 0.0420 respectively).

S4 Fig shows ROC curves for rCBV_max_ at IPE threshold optimization for the three significant experiments after multiple test correction, that is, differentiating proneural from mesenchymal, proneural from neural and proneural from the of subtypes together. Optimal value for distinguishing proneural from mesenchymal was a rCBV_max_ of 3.10 at IPE, whereas for proneural from neural the best cutoff was a rCBV_max_ of 3.01 at IPE. Finally, the optimal threshold for differentiating proneural from the rest of Verhaak subtypes was a rCBV_max_ of 3.12 at IPE.

### 3.3. Overall survival analysis

Table 3 shows Cox proportional hazards regression for rCBV_max_ at IPE and Verhaak subtypes. Vascularity at IPE alone is significatively associated with overall survival. Proneural subtype also yields significant results. When adding subtypes as cofactors, HR, p-values and AUC are maintained relatively stable for every subtype except proneural. In the latter, p-values escalated to non-significance and HR decreased, affecting also confidence intervals. AUC increase is no longer meaningful as neither regressor is significantly associated with survival. All of this points to a blurring in the effect of the biomarker due to the addition of correlated redundant molecular information.

**Table 3.** Cox proportional hazards regression results for a total of 9 models: uniparametric using rCBV_max_ at IPE and Verhaak subtypes and multiparametric using their combination.

|  | rCBV<sub>max</sub> at IPE |  | Verhaak Subtype |  | AUC |
| --- | --- | --- | --- | --- | --- |
|  | HR (CI95) | p-value | HR (CI95) | p-value |  |
| **rCBV<sub>max</sub> at IPE** | 1.81 (1.2, 2.7) | 0.0045* | - | - | 0.5815 |
| **Classical** | - | - | 0.99 (0.5, 2.0) | 0.9748 | 0.5037 |
| **Mesenchymal** | - | - | 0.83 (0.4, 1.6) | 0.5904 | 0.5407 |
| **Neural** | - | - | 0.61 (0.3, 1.3) | 0.1958 | 0.5403 |
| **Proneural** | - | - | 2.58 (1.2, 5.4) | 0.0113* | 0.5847 |
| **rCBV<sub>max</sub>+**<br>**Classical** | 1.81 (1.2, 2.7) | 0.0045* | 0.97 (0.5, 2.0) | 0.9469 | 0.5806 |
| **rCBV<sub>max</sub>+**<br>**Mesenchymal** | 1.87 (1.2, 2.9) | 0.0058* | 1.15 (0.6, 2.4) | 0.7066 | 0.5676 |
| **rCBV<sub>max</sub>+**<br>**Neural** | 1.77 (1.2, 2.7) | 0.0064* | 0.65 (0.3, 1.4) | 0.2741 | 0.5898 |
| **rCBV<sub>max</sub>+**<br>**Proneural** | 1.55 (1.0, 2.5) | 0.0759 | 1.67 (0.7, 4.1) | 0.2577 | 0.6102 |

\* for statistical significance

The same test was performed for rCBV_max_ at ET in the S5 Appendix. As a well-known biomarker, it also shows significant association with overall survival by its own. In this case, however, when adding the subtypes as cofactors, HR and significance is maintained for every subtype, including proneural, sometimes even improving AUC.

## 4. DISCUSSION

In this study, we examined whether the vascular properties of the potentially infiltrated peripheral edema habitat were correlated with Verhaak molecular subclasses, and especially the proneural subtype. The results show that a value of rCBV_max_ higher than 3.12 at IPE is significantly related to proneural subtype.

There are several studies aiming to determine the influence of genetic expression patterns in MRI features within the edema region. Carrillo *et al* pointed out that edema could have prognostic importance in cases when *MGMT* promoter is methylated^30^, which is a common trait in the proneural subtype^31^. Naeini *et al* discovered that T2 and FLAIR volume hyperintensity representing edema was higher in proneural phenotypes^32^ and Zinn *et al* published that, by stratifying into high and low FLAIR radiophenotypes, they could identify glioblastoma subtypes^33^. Finally, the study of MRI perfusion and genetics of GBM from Barajas *et al*, pointed out the need for a deeper understanding of peritumoral non-enhancing tumor for its risk in future progression as its genetic expression pattern differs from that of the enhancing lesion^18^.

In light of these results, in this study we analyzed the radiomic relevance of the edema using a more detailed characterization of edema heterogeneity by differentiating between non-enhancing (i.e. IPE) and vasogenic edema (i.e. VPE), based on ONCOhabitats approach^15^. This allowed to overcome a limitation pointed out in previous studies^31^ and thus identify the IPE as a region with a radiomic relevance when studying the proneural type. In particular, we found a significant association of the rCBV at the IPE with the glioblastoma molecular profile.

The prognosis potential of the rCBV in glioblastoma at the ET ROI has been extensively studied^14,34,35^. The prognosis potential of the rCBV in glioblastoma in the non-enhancing region was also suggested in some studies^15,23,36^. Jain *et al* performed a survival analysis estimating HR for rCBV in both the enhancing and non-enhancing areas adjusting them for the Verhaak molecular subtypes and using the same database as this study^19^. They found that statistical significance for vascularity enhancing areas improved, suggesting additional information provided by molecular profile. However, for the edema region, it did not improve when adding the molecular information. The different molecular behavior of enhancing and non-enhancing regions is consistent with our results. Moreover, our method allowed to find bigger differences when adjusting for the proneural subtype. Thus, the Cox regression model performed in this study suggested that the predictive power of rCBV at non-enhancing areas could be related to its relationship with survival-related mutations.

As Verhaak noted when describing the molecular subclasses, they can be therapeutically relevant^5^. The fact that the biggest differences were found between the proneural and mesenchymal subtypes may point out that their vascular behavior in peritumoral regions varies broadly. As these two have been the most clinically relevant^10,12^, being able to identify them by specific perfusion features can be crucial for diagnosing and treating glioblastoma patients.

Finally, in 2016 the WHO published a glioblastoma classification mainly studying whether the *IDH* gene presents itself as mutant or wild-type^2^. Our findings are in line with the classification: the proneural subtype was the most significantly different subtype from the three remaining and it has proven to be the most closely related to *IDH1* mutations^5^. Unfortunately, not enough information was available to confirm if *IDH*-mutant vascularity at the peripheral areas was significantly different from wild-type.

There were some limitations in this study. Firstly, though we were able to correlate the IPE habitat and the Verhaak subclasses, due to the retrospective nature of the study, we were not able to standardize MRI acquisition protocols. Secondly, it can be difficult to correctly asses the exact location of the infiltrated edema by a noninvasive manner. These could affect the potential relationships with molecular markers. Nonetheless determining the IPE habitat by an automated method for calculating the maximum of the cerebral blood volume to perform seems robust enough for our purpose, as shown by Álvarez-Torres *et al*^23^. Finally, despite having significant results, the sample size available in the dataset may be a statistical limitation.

Our study relies on the use of an automatic procedure to determine a more precise peritumoral ROI based on an open service^24^ and the results can be replicated using the TCGA-GBM open dataset^26^.

## 5. CONCLUSIONS

In conclusion, high IPE vascularity features are associated with the proneural subtype. Global vascularity differences between the four subtypes exist in this region especially due to proneural and mesenchymal influence. rCBV_max_ at IPE is related to overall survival and carries specific molecular information.

## ACKNOWLEDGMENTS

This work was partially supported by: MTS4up project (National Plan for Scientific and Technical Research and Innovation 2013-2016, No. DPI2016-80054-R) (JMGG); H2020-SC1-2016-CNECT Project (No. 727560) (JMGG), Fundació Bancaria laCaixa (LCF/TR/CI16/10010016) and H2020-SC1-BHC-2018-2020 (No. 825750) (JMGG). M.A.T was supported by DPI2016-80054-R (*Programa Estatal de Promoción del Talento y su Empleabilidad en I+D+i*). EFG was supported by the European Union’s Horizon 2020 research and innovation program under the Marie Skłodowska-Curie grant agreement No 844646. We gratefully acknowledge the COST Association for its CA18206 - Glioma MR Imaging 2.0 European project which supports the research on glioblastoma.

## SUPPORTING INFORMATION CAPTIONS

**S1 Table. Clinical data of the final cohort.** All clinical data is obtained from the TCGA-GBM open database^26^. Subtype information is retrieved from the UCSC Xena platform compilation^28^ which is based on Brennan *et al*^27^.

**S2 Table. rCBV**_**max**_, **rCBV**_**mean**_ **and rCBV**_**median**_ **at every habitat for each subject.**

**S3 Appendix. Mann Whitney U-test comparing rCBV**_**max**_ **at each habitat for Verhaak subtypes.**

**S4 Fig. ROC curves for rCBV**_**max**_ **at IPE threshold optimization for significant experiments.** Significant experiments: distinguishing proneural from mesenchymal, proneural from neural and proneural from the rest.

**S5 Appendix. Cox proportional hazards regression for rCBV**_**max**_ **at ET and Verhaak subtypes.**

